# Effects of topical PaiTeLing in nude mice implanted with human condyloma acuminatum tissue infected with HPV 6, 31, and 81: comparison with imiquimod and interferon-α-2b

**DOI:** 10.1101/563536

**Authors:** Rong Xu, Li Wang, Jianmei Hou, Jun Li, Zhiyan Fan, Liangcai Wu, Congzhong Zhu, Miaomiao Ma, Huiping Wang, Shuping Hou

## Abstract

**BACKGROUND:** The standard treatment for condyloma acuminatum is topical imiquimod. In the current study, we used a mouse model to compare the effects of an herbal medication PaiTeLing.

**METHODS:** Lesion tissue was obtained from a woman with genital condyloma acuminatum. DNA genotyping revealed HPV6, 31, and 81. Tissue prism (0.5 cm^3^) was implanted to BALB/C nude mice, 22 days after the implantation, mice began to receive topical treatment with imiquimod, interferon-α-2b gel, or PaiTeLing over the site of implantation for 2 consecutive weeks. Mice receiving tissue implantation but no other intervention was included as a control. Skin tissue was collected for H&E staining and anti-CD207 immunohistochemistry. Blood was collected to determine a panel of cytokines.

**RESULTS:** H&E staining showed lower number of koilocytes and higher number of Langerhans cells in the treatment groups, particularly in mice receiving imiquimod or PaiTeLing. Blood levels of TNFα, IL-2, INF-γ and IL-12p70 were increased in the treatment groups, particularly in mice receiving imiquimod or PaiTeLing.

**CONCLUSION:** Immune response in nude mice infected with HPV6, 31, and 81 is enhanced by treatment with imiquimod, interferon-α-2b and PaiTeLing. Effects of imiquimod and PaiTeLing seems to be stronger than interferon-α-2b.

## INTRODUCTION

Condyloma acuminatum (CA) is a wide-spread sexually transmitted disease with increasing prevalence during the past decades [1]. CA is caused by human papillomavirus (HPV) infection. Infection with high-risk HPV strains is closely associated cervical cancer as well as other type of cancer in the anogenital region [2, 3]. HPV16 is a major risk for the development of oropharyngeal squamous-cell-carcinoma [4, 5]. HPV infection causes mental distresses and produces negative impact on sexual life and social well-being of infected individuals. Clearance of HPV virus is the goal for the treatment of genital warts and could prevent cancers. Most of the currently available treatments focus on clearing the externally visible warts rather than targeting the viruses [6,7]. An animal model for CA that look into both the local and systemic immune responses is essential for development of new treatments [8,9].

Host immune responses and chronic inflammation have been shown to be important risk factors for HPV-related carcinogenesis [10]. Increased duration of HPV infection could influence disease development, indicating the importance of the host immune response to HPV clearance and HPV-related carcinogenesis. Immune responses to invading HPV consist of innate and adaptive immune systems. Langerhans cells are immature dendritic cell (mDC), and the only professional antigen-presenting cells in the epidermis. [11,12] HPV infection results in a net loss of Langerhans cells at the site of infection. [13] HPV also interferes with antigen presentation and processing machinery in Langerhans cells, [14, 15] and alters chemokine and cytokine expression by LC. [16, 17] Cytokines TNF-α, IL-2, IFN-γ and IL-12p70 are associated with HPV clearance. [18–21] More than 90% of cervical squamous cancers and 75% of cervical lesions with koilocytes harbor HPV DNA [21]. As a result, koilocytes in CA may be associated with HPV infection.

Imiquimod is an immuno-modulating agent with actions at multiple levels of the adaptive immune system. Imiquimod activates the cells of the immune system via toll like receptor, and increase the secretion of cytokines, including TNF-α, IL-1β, and IL-6. [22] PaiTeLing (Beijing Paitborn Company, the Chinese Academy of Medical Sciences) is an herbal medication widely used to treat genital warts in China. The main components are honeysuckle and fructus cnidii, Indigowoad Leaf, sophorae flavescentis, Brucea javanica, and Hedyotis diffusa. [23] A previous study suggested that PaiTeLing could promote HPV clearance and regression of cervical lesions. [24] PaiTeLing selectively destroys cancer cell membranes, including cytoplasmic and mitochondrial membranes, thereby suppressing cancer cell proliferation. In addition, PaiTeLing could also destroy viruses that reside inside the cells [25]. Interferons are a group of natural messenger proteins with antiviral actions. Systemic administration of interferons after ablative treatment for anogenital warts has been advocated to increase clearance and decrease recurrence rate. [26] Human interferons are a class of small proteins and glycoprotein cytokines (15-28 kD) produced by T cells, fibroblasts, and other cells in response to viral infection and other biologic stimuli. Interferons bind to specific receptors on cell membranes to produce a variety of biological actions, include inducing enzymes, suppressing cell proliferation, inhibiting viral proliferation, enhancing the phagocytic activity of macrophages, and augmenting the cytotoxic activity of T lymphocytes.[27] In the current study, we used a mouse model of human CA to examine the potential effects of topical imiquimod, PaiTeLing and interferon-α-2b.

## MATERIALS AND METHODS

### 1.1 Experimental animals

Female BALB/C-nu nude mice (4-6 weeks of age; Experimental Animal Center, the Chinese Academy of Medical Sciences Hematology Hospital and Institute of Hematology) were used in this study. The mice were housed in a specific-pathogen free facility.

### 1.2 Human condyloma acuminatum tissue and genotyping

Human condyloma acuminatum tissue was obtained from an elderly woman with informed consent prior to any treatment. The diagnosis was based on acetic acid white test and pathological examination. HPV genotyping was carried out using a 23-type polymerase chain reaction-based kit from Yanneng Biotechnology (YZB/County 3881-2014). The results showed HPV6, HPV31 and HPV81.

### 1.3 Tissue implantation

The wart tissue was removed under aseptic condition, transferred into EP tubes containing Ringer lactate solution, and soaked in a solution containing 1-ml D-hanks solution and 2.5-ml streptomycin/penicillin for 45 minutes prior to slicing into 0.5-cm^3^ tissue blocks. Mice were anesthetized with sodium pentobarbital (50 mg/kg; intraperitoneally), and implanted with condyloma acuminatum tissue under aseptic condition through an incision to beneath the skin of the right dorsal. The incision was closed layer by layer using 5-0 silk suture. The surgery site was covered with sterile vaseline gauze and dressed with elastic bandage for 24-48 hours. Povidone iodine was used to clean the wound for about one week.

### 1.4 Treatment

Starting from 3 weeks after the implantation, the incision site was treated with topical imiquimod (5%) once every other day, PaiTeLing (1:20 or 1:2) or interferon-α-2b twice a daily. The treatment lasted for 2 weeks.

### 1.5 Hematoxylin and eosin stain

Upon the completion of the treatment, mice were sacrificed by cervical dislocation. The skin at the incision site was fixed in 4% paraformaldehyde for 48-h at room temperature, embedded in paraffin wax. Paraffin was removed from tissue slices (4 μm) using xylene and rehydrated by brief submersion in a series of aqueous alcohol solutions of decreasing alcohol concentration prior to standard staining with hematoxylin and eosin (HE) and examination under light microscopy.

### 1.6 Immunohistochemical analysis

Immunohistochemical analysis was performed in formalin-fixed paraffin-embedded tissues (FFPET). After deparaffinization and dehydration, heat-induced antigen retrieval was carried out. The sections were incubated overnight with a primary antibody against CD207 (anti-rabbit polyclonal; R&D NOVUS made in Colorado; identification of product NB100-56733SS), a marker for mDC expressing Langerin, at 1:500 dilution [28]. The secondary antibody was EnVision®+Dual. Five randomly selected fields for each sample were included in the cell counting and analysis.

### 1.7 Cytokine analysis

MSD proinflammatory Panel I (MesoScale Discovery, Gaithersburg, MD, USA), a highly sensitive multiplex enzyme-linked immunosorbent assay (ELISA), was used to examine serum concentration of 10 cytokines, including interferon γ (IFN-γ), interleukin (IL)-1β, IL-2, IL-4, IL-6, IL-8, IL-10, IL-12p70, IL-13, and tumor necrosis factor α (TNF-α), as described previously [29]. The plates were run on a Meso Quickplex machine; data were analyzed using the MSD Discovery Workbench software v4.0.

### 1.8 Statistical analysis

All statistical analyses were performed using the SPSS 16.0 software. Data were analyzed with one-way analysis of variance (ANOVA), and presented as mean ± standard deviation. Statistical significance was set at P<0.05.

## Results

### Skin tissues at the incision site

Mice receiving human condyloma acuminatum tissue was characterized by typical features, including koilocytes. In contrast to the absence of koilocytes in normal skin tissue, skin at the site of incision in mice receiving human condyloma acuminatum tissue but no other treatment contained many koilocytes (Figure 2a/b). The number of koilocytes was significantly reduced by imiquimod, interferon-α-2b and PaiTeLing (1:20 and 1:2), with seemingly more robust effects for imiquimod and PaiTeLing (Figure 2c/e). A quantitative analysis showed that the 1:2 PaiTeLing group had the largest reduction in koilocytes, followed by imiquimod, 1:20 PaiTeLing, and then interferon-α-2b (Figure 2g).

### Cytokines

Among the 10 factors examined using the MSD proinflammatory Panel I, IFN-γ, IL-2, IL-12p70 and TNF-α play important roles in the microenvironment of CA. Therefore, we analyzed the levels of these four cytokines.

Levels of four cytokines were significantly higher in all 4 groups of mice receiving treatments vs. the negative control (Figure 1a/c/e/g). Mice receiving mice receiving 1:20 PaiTeLing group and 1:2 PaiTeLing group had lower TNF-α levels, compared with mice receiving imiquimod group (P<0.001, P<0.01, respectively) (Figure 1h), but there was no statistical significance compared with the interferon-α-2b group (Figure 1h). Mice receiving imiquimod group and interferon-α-2b group had higher IL-2 levels than 1:20 PaiTeLing group (P<0.001, P<0.01, respectively), but there was no statistical significance compared with the 1:2 PaiTeLing group (Figure 1d). All four groups of mice receiving treatments had similar IL-12p70 levels (Figure 1f). In comparison with the 1:20 PaiTeLing group and the 1:2 PaiTeLing group, mice receiving imiquimod (P<0.01, P<0.01, respectively) and mice receiving interferon-α-2b (P<0.01, P<0.01, respectively) had lower IFN-γ levels (Figure 1b).

**Fig 1:**
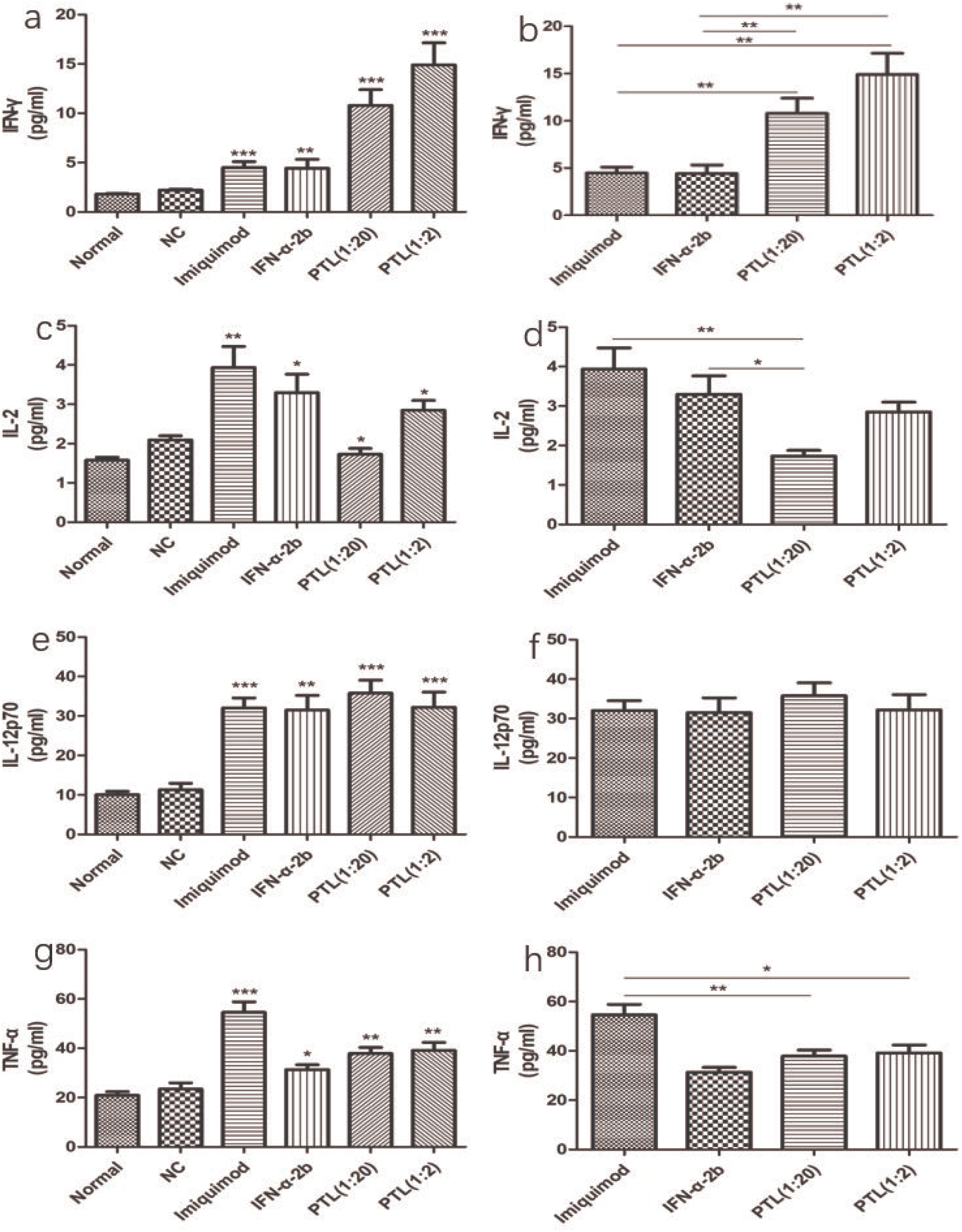
The level of blood cytokines. a-b: interferon-γ (IFN-γ); c-d: interleukin-2 (IL-2); e-f: interleukin-12p70 (IL-12p70); g-h: tumor necrosis factor-α (TNF-α); Imiquimod, interferon-α-2b, and PaiTeLing increased the levels of IFN-γ, IL-2, TNF-α, and IL-12p70. Effects of PaiTeLing and imiquimod were more pronounced than interferon-α-2b. Error bars: SD. * P<0·05; ** P<0·01; *** P<0·001.

**Fig 2:**
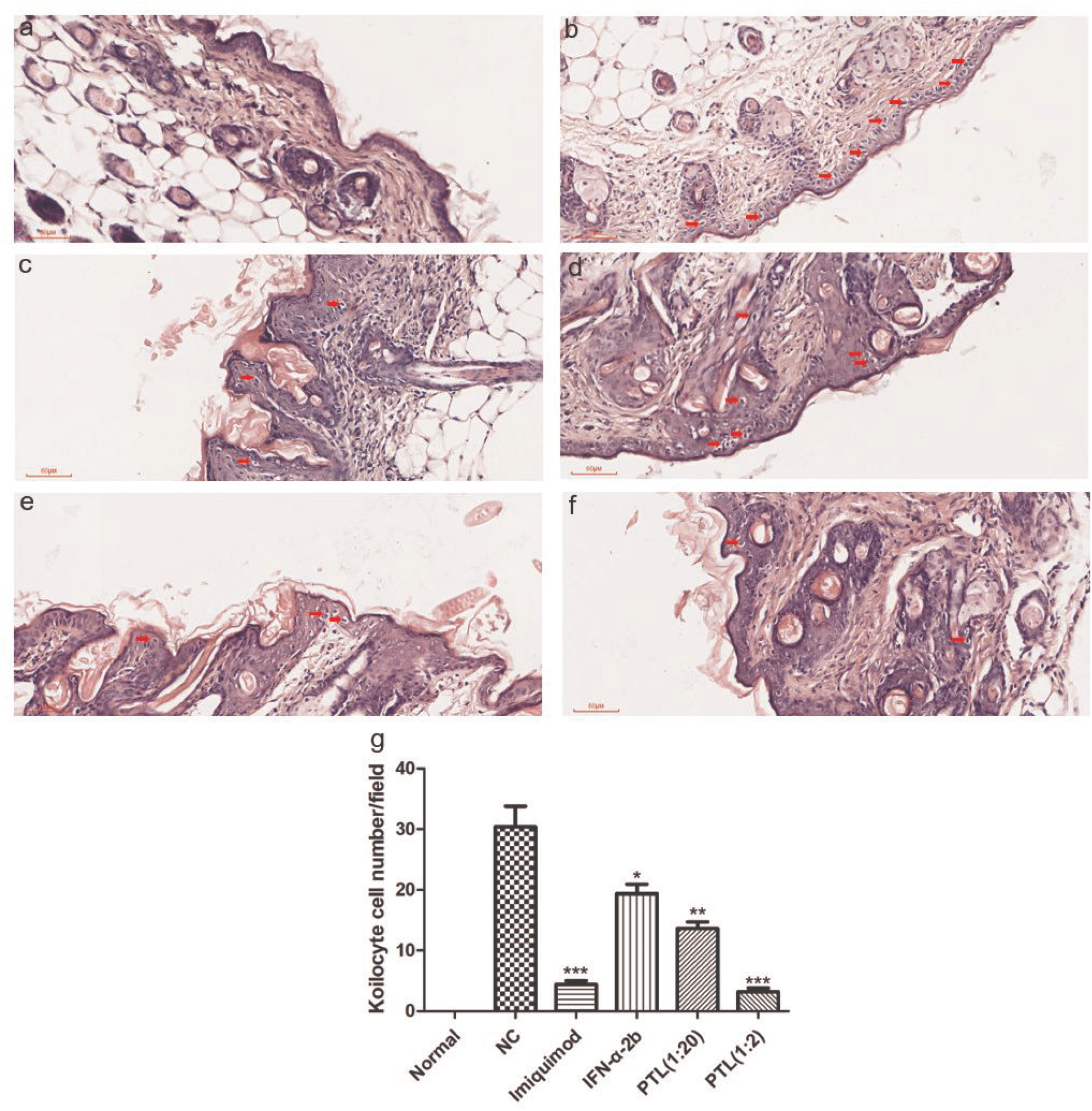
Skin tissue at the incision site. a: healthy control; b: mice receiving human CA tissue but no other treatment; c: imiquimod; d: interferon-α-2b; e-f: 1:20 and 1:2 PaiTeLing. Arrows: koilocytes. Staining with hematoxylin and eosin (HE); magnification: 200; g: Summary statistics. Error bars: SD. * P<0·05; ** P<0·01; *** P<0·001.

### Immunohistochemical staining for LC

LC was observed in all the slides(Figure 3a-f). The number of LC in the normal group was lower than in mice receiving human condyloma acuminatum tissue without any treatment (5.33±0.94 vs. 6.33±1.25; Figure 3g). The number of LC in the four treatment groups was significantly higher than in the untreated control group (Figure 3g). The number of LC was highest in the imiquimod group (28 ±1.70), and lowest in the interferon-α-2b group (14 ± 1.70). The number of LC in the 1:2 and 1:20 PaiTeLing groups were higher than in the interferon-α-2b (24 ±1.63, 21 ±1.25) (P<0.01, P<0.01, respectively) (Figure 3h). The number of LC in the imiquimod groups was higher than in the 1:20 PaiTeLing group (24±1.63, 21 ±1.25) (P<0.01, P<0.01, respectively) (Figure 3h), but there was no statistical significance compared with the 1:2 PaiTeLing group (Figure 1d).

**Fig 3:**
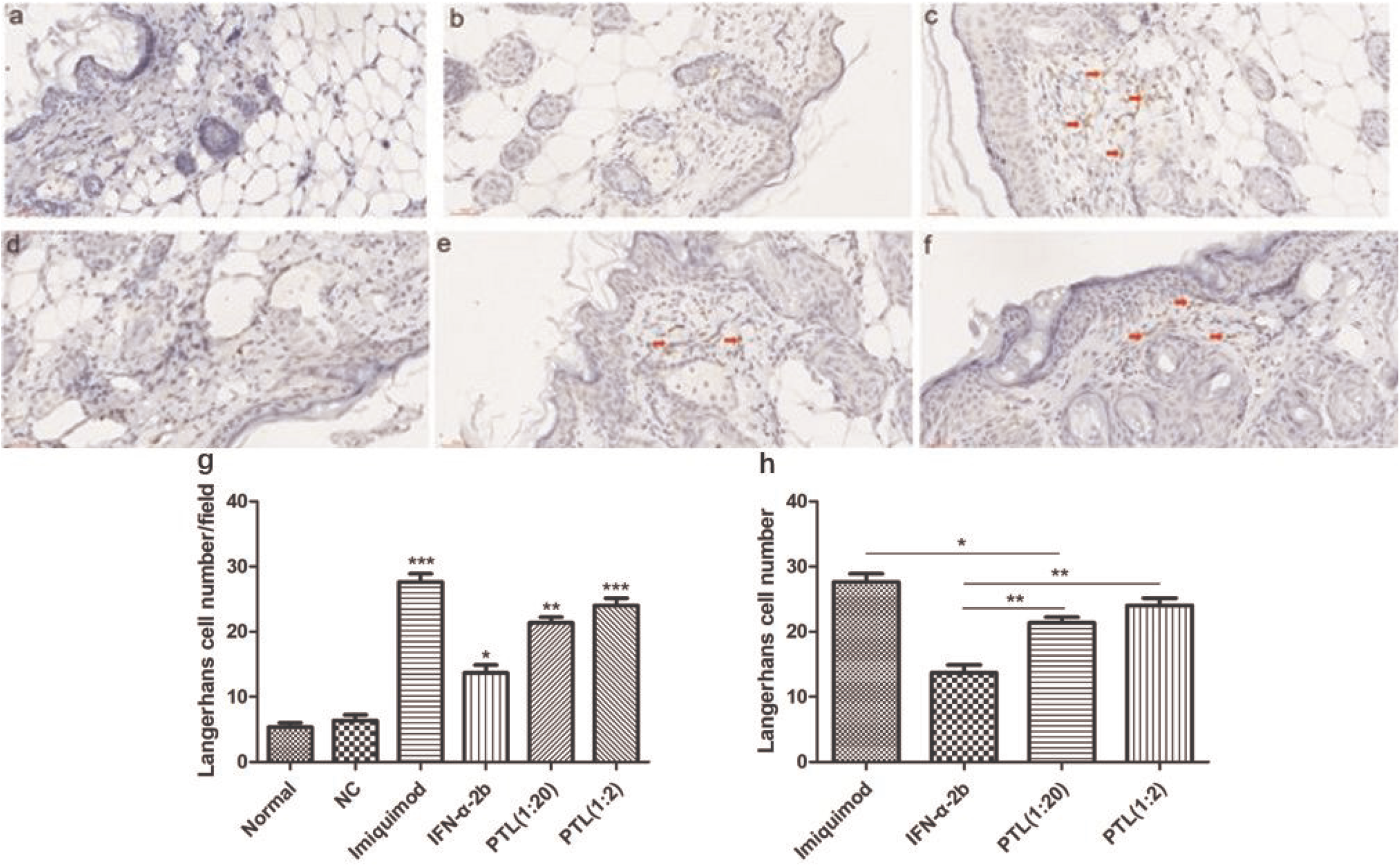
Immunohistochemical staining for CD207. a: healthy control; b: mcie receiving human CA tissue but no other treatments; c: imiquimod; d: interferon-α-2b; e-f: 1:20 and 1:2 PaiTeLing. Arrows: Langerhans cells; magnification: 200; g-h: Summary statistics of the number of Langerhans cells. Asterisks (*): Langerhans cells. Error bars: SD. * P<0·05; ** P<0·01; *** P<0·001.

## Discussion

HPV infection is an important factor in the development of CA.[30] Analyzing the immune responses to HPV infection is useful to determine the developmental mechanism of CA. Cutaneous infections caused by HPV are usually recurrent and are among the most troublesome conditions presenting to dermatologists.[31] Common warts are the most frequent manifestation to skin HPV infection. Warts are usually self-limiting but spontaneous resolution may take months to years. Spontaneous clearance rate has been reported to be 23% at 2 months, 30% at 3 months and 65-78% at 2 years.[32] Clearance of HPV infection is the ultimate goal for the treatments.[33] Strong evidence suggests that HPV specific cell-mediated immune (CMI) response is pivotal in clearing HPV.[34] An increased understanding of the importance of cellular immunity in clearance of HPV infections has led to therapeutic trials of interferons [35–37] and imiquimod [38, 39] in patients with anogenital condylomas. PaiTeLing is widely used to treat CA to remove genital warts and to prevent recurrence.

The number of koilocytes reflect HPV infection. In the current study, the number of koilocytes at the skin incision site as significantly higher than in the normal control (not receiving human CA tissue), indicating successful infection of the mice. Topical treatment with all 4 medications in the current study decreased the number of koilocytes at the incision site. The order of the effect magnitude was 1:2 PaiTeLing > imiquimod > 1:20 PaiTeLing > interferon-α-2b, suggesting that PaiTeLing is an effective agent.

T helper type 1 (Th1) response is important for the clearance of HPV infection; conversely, lack of Th1 response has been associated with persistent infection and HPV-associated neoplasia.[34] Th1 cells produce IFN-γ, TNF-α, and IL-2, and are critical for cellular immune responses.[40] Accessory cell-derived cytokines, such as IL-12, promote Th1 cell development, and are also important regulators of T-cell functions.[34] LCs are antigen-presenting cells of the skin, and represent the first link of T cell immune responses.[41] In the current study, the levels of IFN-γ, TNF-α, IL-2 and IL-12p70 in mice receiving human CA tissue were slightly higher than that in the healthy control, and were significantly increased by all 4 treatment, indicating that potentiation of the immune response. Specifically, TNF-α and IL-2 were the highest in the imiquimod group, IFNγ was highest in the 1:2 PaiTeLing group, IL-12p70 was highest in the 1:20 PaiTeLing group. IL-2 was higher in the interferon-α-2b group than the 1:2 PaiTeLing group. The number of LCs was highest in the imiquimod-treated group, and lowest in the interferon-α-2b group.

In summary, the results from the current study suggest that topical treatments with imiquimod, interferon-α-2b and PaiTeLing could help to promote host immune response to HPV. Effects seem to be strongest with imiquimod, followed by PaiTeLing and then interferon-α-2b. Considering the side effects of imiquimod (e.g., excoriation, erosion, ulceration, burning sensation, pain, itching and hypopigmentation,[42] PaiTeLing is a viable alternative.

